# Network regularized Accelerated Failure Time Models for Robust Biomarker Identification

**DOI:** 10.1101/2025.05.30.657087

**Authors:** Yang Liu, Jason Huse, Kasthuri Kannan

## Abstract

**Background:** High-dimensional genomic studies in cancer research face significant challenges when analyzing survival outcomes, particularly in p ≫ n scenarios where thousands of genes are measured across relatively small patient cohorts. Traditional penalized regression methods like lasso and ridge regression apply uniform shrinkage without considering underlying biological relationships between genes, often resulting in unstable feature selection and reduced interpretability. Network-regularized methods that incorporate prior biological knowledge have emerged as promising alternatives, yet systematic comparisons of different network penalties in accelerated failure time (AFT) models remain limited.

**Methods:** We developed and evaluated Weibull AFT models incorporating two network-regularized approaches: inverse-degree penalties that apply stronger shrinkage to highly connected hub genes, and graph Laplacian penalties that promote smoothness across connected nodes. We conducted comprehensive simulations across varying signal strengths, censoring rates (30%, 50%, 80%), and noise levels, followed by real-data applications on three cancer datasets (TCGA-KIRC, TCGA-COAD, and IDH-wildtype gliomas) using both STRING protein-protein interaction (PPI) networks and co-expression networks. Performance was assessed using estimation and prediction mean squared error, false positive/negative rates, concordance index, and pathway enrichment analysis.

**Results:** The inverse-degree penalty demonstrated consistently more conservative feature selection behavior, achieving substantially lower false positive rates while maintaining competitive predictive performance compared to Laplacian penalties. In simulation studies, the inverse-degree method achieved false positive rates of 0.126 versus 0.299 for Laplacian penalties under strong signal conditions. Real-data applications showed comparable predictive accuracy (C-index: 0.61-0.73) between methods across cancer types. Both methods successfully identified biologically relevant pathways including mTORC1 regulation and fatty acid metabolism in kidney renal clear cell carcinoma (KIRC), and IL-17/chemokine signaling in IDH-wildtype gliomas. Notably, in colorectal adenocarcinoma (COAD) analysis, the inverse-degree method identified ten coherent pathways centered on inflammation, cell-cycle regulation, and proteasome function, while the Laplacian approach introduced potentially spurious neurotransmitter-release associations, demonstrating the inverse-degree penalty’s superior biological interpretability.

**Conclusions:** Network-regularized AFT models with inverse-degree penalties offer a valuable tool for biomarker discovery in high-dimensional genomic survival analysis, providing conservative feature selection and superior biological interpretability while maintaining competitive predictive performance. The accompanying DegreeAFT R package makes these methods accessible to the broader research community, facilitating adoption in precision oncology applications where identifying reliable prognostic signatures is critical for treatment decision-making.

## 1 Introduction

High-dimensional genomic studies in cancer research commonly involve measuring tens of thousands of genes across relatively small patient cohorts, creating challenging p≫ n scenarios where the number of covariates vastly exceeds the sample size (Goeman, 2010). This dimensional imbalance, coupled with complex multicollinearity among genomic features, poses significant challenges for traditional survival analysis methods. Classical penalized regression approaches, such as the lasso (Tibshirani, 1996) and ridge regression (Hoerl & Kennard, 1970), apply uniform shrinkage across all coefficients without considering the underlying biological relationships between genes, often resulting in unstable feature selection and reduced interpretability of identified biomarkers (Zou & Hastie, 2005).

The biological reality of gene regulation presents additional complexity that standard regularization techniques fail to address. Genes do not operate independently but are organized within intricate regulatory networks, including co-expression modules, protein-protein interaction (PPI) networks, metabolic pathways, and transcriptional regulatory circuits (Barabási & Oltvai, 2004).These structured dependencies among genomic features violate the independence assumptions of conventional penalized methods, often leading to reduced prediction accuracy and limited biological interpretability.

To overcome these challenges, researchers have developed network-regularized regression methods that incorporate prior biological knowledge into the modeling framework. These approaches use known gene-gene interactions to guide the shrinkage process, enhancing both statistical performance and the biological relevance of models. Several strategies have emerged for integrating network information into penalized regression. In 2005, Tibshirani et al. introduced the fused lasso, which augments the classic lasso with an L1 penalty on differences between coefficients of connected predictors, thereby enforcing both sparsity and smoothness in the estimated effects. Building on this, Li & Li, 2008 developed a network-constrained regularization framework for genomic data that incorporates known gene–gene interactions into the penalty structure. Their approach uses a quadratic graph Laplacian penalty to promote local consistency and stability across connected genes and further extends the elastic net by integrating network-derived weights, striking a balance between grouping correlated variables and maintaining overall sparsity. Network-based group penalties treat connected components or functional modules as groups, using group Lasso or group Ridge penalties to promote coordinated selection of biologically related genes (Zhu et al., 2009). These methods have shown promise in genomic applications where biological pathways and gene networks provide natural grouping structures.

Among these approaches, two strategies have received particular attention in survival analysis applications. The graph Laplacian penalty has been studied for its effectiveness in promoting smooth coefficient profiles across connected genes in biological networks. The inverse-degree penalty represents an alternative strategy that applies node-specific ridge penalties inversely proportional to each gene’s connectivity (degree) within the network. This approach recognizes that highly connected hub genes may be more prone to false positive selection due to their extensive interactions and therefore applies stronger shrinkage to these nodes. Veríssimo et al. (2016) demonstrated the utility of inverse-degree penalties in Cox proportional hazards models, showing improved variable selection and prognostic performance in high-dimensional genomic datasets. However, their application to AFT models has not been systematically investigated.

Despite the growing interest in network-regularized survival models, several critical gaps remain in our understanding of their relative performance and optimal application conditions. While both Laplacian and inverse-degree penalties have shown promise in isolated studies, there has been no systematic comparison of their performance under realistic conditions for high-dimensional survival analysis, particularly in the context of AFT models. Existing evaluations often rely on simplified simulation scenarios that may not capture the complexity of real genomic data and survival outcomes (Simon et al., 2011). The impact of network structure characteristics on penalty performance remains poorly understood, as real biological networks exhibit diverse topological properties, including modular organization with varying degrees of overlap between functional modules (Fortunato, 2010). The performance of different network penalties may vary substantially depending on whether the underlying network exhibits clear, non-overlapping modules or more complex, overlapping community structures that better reflect biological reality.

Furthermore, survival data in clinical studies are subject to various sources of noise and complexity, including censoring patterns, measurement error, and heterogeneous patient populations (Klein & Moeschberger, 2003). The robustness of network-regularized methods under these realistic conditions has not been thoroughly evaluated, particularly for AFT models where the relationship between covariates and survival times is assumed to be multiplicative rather than proportional (Wei, 1992). AFT models offer several advantages over Cox proportional hazards models, including more straightforward interpretation of covariate effects and greater flexibility in modeling survival distributions, yet they have received less attention in the network-regularized survival analysis literature.

To address these knowledge gaps, we propose and evaluate a Weibull AFT model incorporating inverse-degree penalties for high-dimensional survival analysis. Building upon the simulation framework established by Angelini et al. (2024), we conduct comprehensive comparisons between inverse-degree and Laplacian penalties under biologically plausible conditions. Our evaluation considers multiple network structures representing different levels of modular organization, varying censoring rates, and different noise levels to provide practical guidance for method selection in genomic survival studies. The primary contributions of this work include the first systematic comparison of inverse-degree versus Laplacian penalties in the context of high-dimensional AFT models, evaluation of penalty performance across diverse network topologies and survival data characteristics, and practical recommendations for selecting appropriate network-regularized methods based on study-specific conditions and objectives. These findings will help inform the selection and application of network-regularized survival models in genomic biomarker discovery and prognostic modeling.

## 2 Methods

### 2.1 Data Collection and Preprocessing

Gene expression and clinical data were obtained from The Cancer Genome Atlas (TCGA) or the Glioma Longitudinal Analysis Consortium (GLASS) dataset. The datasets analyzed included TCGA-KIRC, TCGA-COAD and IDH-wildtype gliomas. These datasets were included to assess model performance across cancer types. Prior to analysis, genes with more than 80% zero expression values across samples were removed, and rows with entirely missing values were excluded. Initial gene screening was performed using log-rank tests, retaining the top 500 genes or statistically significant genes with the lowest nominal p-values when FDR-adjusted p-values yielded no significant results. Samples were aligned between expression and survival data matrices to ensure consistency.

### 2.2 AFT Model

The AFT model assumes covariates act multiplicatively on the survival time scale. For subject *i* with survival time *T*_*i*_ and covariate vector *x*_*i*_ ∈ ℝ^*p*^, the model is defined as:

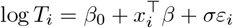

Where *β*_0_ is the intercept, *β* is the vector of regression coefficients, *σ >* 0 is the scale parameter, and *ε*_*i*_ follows a standard extreme value distribution. Under the Weibull distribution assumption, the scale parameter is 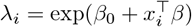 and the shape parameter is *k* = 1*/σ*. The probability density function (PDF) and survival function for subject *i* are:

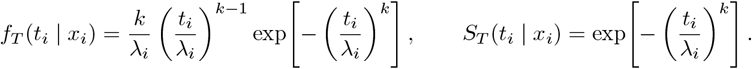

### 2.3 Network-Regularized AFT Model

The objective function incorporates L1 (lasso) and network-based penalties:

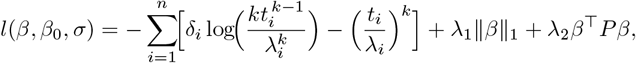

where *δ*_*i*_ ∈ {0, 1} is the event indicator, *λ*_1_ ≥ 0 controls sparsity, *λ*_2_ ≥ 0 controls network regularization, and *P* is either the Laplacian matrix *L* or the inverse-degree matrix *D*^−1^. In log-time coordinates with *y*_*i*_ = log *t*_*i*_ and rescaled residuals 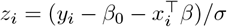, the gradients are:

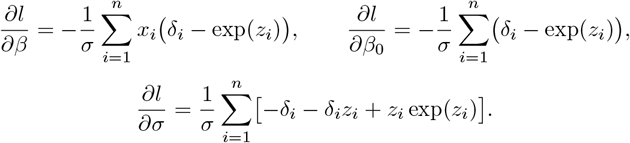

The non-smooth L1 penalty is handled using proximal gradient descent, alternating between gradient steps for smooth components and proximal operators for the L1 term:

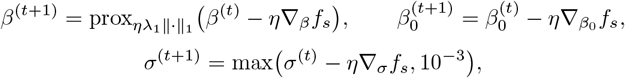

where *f*_*s*_ is the smooth part of the objective function, *η* = 1*/L* is the step size, and 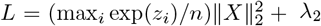 max(diag(P)) is the Lipschitz constant. The detailed derivations is described in Appendix.

### 2.4 Simulation Study

Simulations were conducted over 100 replicates following established frameworks for network-constrained survival analysis, evaluating two network structures: star-shaped subnetworks with each transcription factor (TF) connected to 10 unique target genes, and complex structures with genes regulated by multiple TFs, forming overlapping regulatory modules. For each scenario, sample sizes were set to *n* = 165 (weak signal) or *n* = 413 (strong signal), with dimensions *p* = 220 (weak) or *p* = 1,100 (strong) features, including 20 TFs or 100 TFs. Active coefficients included 88 non-zero values, with *β*∈ [0.5, 0.1] (positive) or *β*∈ [−0.5, − 0.1] (negative), and a correlation of *ρ* = 0.7 between TFs and target genes. Censoring rates were 30%, 50%, and 80%, with noise levels *σ* ∈ {0.5, 1.0, 1.5}. Survival times were generated from a Weibull distribution with shape *k* = 1*/σ* and scale 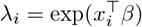. Censoring was applied randomly. Adjacency matrices reflected TF-gene regulatory relationships, with degree matrices computed as *D*_*ii*_ = ∑_*j*_ *A*_*ij*_. The inverse-degree matrix was scaled by mean(rowSums(|*L*|))*/* mean(diag(*D*^−1^)) to ensure comparable penalty magnitudes. Model performance was evaluated using estimation mean squared error mean((*β*′ − *β*)^2^), prediction mean squared error mean((*Xβ*′ − *Xβ*)^2^), false negative rate 1 − |selected ∩ active|*/*|active|, false positive rate |selected ∩ inactive|*/*|inactive|, and network selection rate |selected|*/p*. Hyperparameters *λ*_1_ ∈ {0.005, 0.01, 0.02} and *λ*_2_ ∈ {0.005, 0.01, 0.025, 0.05} were selected via 5-fold cross-validation, optimizing the concordance index (C-index).

### 2.5 Real Data Analysis

Biological networks were constructed using two approaches. STRING PPI networks (v12.0) were filtered for high-confidence interactions (combined score ≥ 200) among genes in the expression datasets. WGCNA co-expression networks were built using the WGCNA package with unsigned correlation networks, selecting soft-thresholding power to achieve scale-free topology (*R*^2^ ≥ 0.8). Genes were pre-screened using log-rank tests (median split), retaining the top selected genes with the lowest nominal *p*-values when FDR-adjusted *p*-values were non-significant. Networks were restricted to these genes. Models with inverse-degree and Laplacian penalties were fitted using the proximal gradient algorithm. Evaluation included 5-fold cross-validation for hyperparameter tuning, performance assessment via C-index and integrated AUC (iAUC), feature standardization within each cross-validation fold to prevent data leakage, and analysis of selected genes for biological relevance.

### 2.6 Pathway Enrichment

To investigate the biological relevance of genes identified by our network-regularized AFT models, we performed pathway enrichment analysis on three cancer datasets: IDH-wildtype glioma, KIRC, and COAD. This analysis aimed to uncover the underlying biological pathways associated with survival outcomes, leveraging the gene sets derived from three distinct approaches: differentially expressed genes (DEG), STRING protein-protein interaction networks, and WGCNA.

For each dataset and method, genes were ranked based on their regression coefficients from the AFT models, reflecting their association with survival time. We selected the top 20% and bottom 20% of genes, those with the strongest positive and negative impacts on survival, respectively for pathway enrichment. Enrichment analysis was conducted using ToppGene with pathways considered significantly enriched if they met a false discovery rate (FDR) threshold of less than 0.05.

## 3 Results

### 3.1 Simulation Study Results

The simulation study evaluated the performance of the proposed Weibull AFT model with inverse-degree (MAIN) and Laplacian (REF) penalties across varying conditions of signal strength, censoring rates, and noise levels. Performance metrics included estimation mean squared error (EMSE), prediction mean squared error (PMSE), false negative rate (FNR), false positive rate (FPR), and network selection rate (NSR).

The results demonstrate that the proposed inverse-degree penalty method is more conservative in feature selection, consistently achieving lower false positive rates and network selection rates compared to the Laplacian penalty approach. Also, the behavior of the two penalties is remarkably similar in the overlapping and non-overlapping networks. In the strongest-signal, lowest-noise scenario (strong signal, 30 % censoring, *σ* = 0.5) the inverse-degree penalty (MAIN) reduces the false-positive rate (FPR) from 0.299 to 0.126 in the overlapping topology and from 0.289 to 0.069 in the non-overlapping topology, while the Laplacian reference (REF) retains a much lower false-negative rate (FNR, 0.035 versus 0.430 in the overlap case and 0.039 versus 0.445 in the non-overlap case). The magnitude of these gaps changes only marginally as noise increases or censoring becomes more severe, indicating that network overlap has little practical effect on the trade-off between sparsity and smoothness.

EMSE and PMSE are small in absolute terms and differ only modestly between the two penalties across all scenarios. Both penalties show the expected monotone rise in PMSE as *σ* increases and a gradual deterioration as censoring intensifies, yet MAIN consistently achieves the lower FPR.

### 3.2 Real Data Application Results

For KIRC dataset, we obtained 731 genes with significant p-values of log rank test and we use top 500 genes of COAD and glioma dataset for network construction since no gene is significant after multiple tests correction. The real data application assessed the performance of both penalty methods on three cancer datasets using STRING and WGCNA networks, as shown in Table 3. The selected gene markers are listed in Supplementary Table 1. Performance was evaluated via the C-index, iAUC, the number of genes selected, and computational time.

**Table 1.** Overlap-simulation benchmark (100 replicates, 5-fold CV)

| Choice | Censor | $\sigma$ | EMSE | | PMSE | | FNR | | FPR | | NSR | |
| --- | --- | --- | --- | --- | --- | --- | --- | --- | --- | --- | --- | --- |
|  |  |  | MAIN | REF | MAIN | REF | MAIN | REF | MAIN | REF | MAIN | REF |
| weak | 0.3 | 0.5 | 0.029 | 0.015 | 0.308 | 0.176 | 0.309 | 0.062 | 0.471 | 0.546 | 0.559 | 0.703 |
| weak | 0.3 | 1 | 0.038 | 0.023 | 0.476 | 0.330 | 0.359 | 0.137 | 0.488 | 0.585 | 0.549 | 0.696 |
| weak | 0.3 | 1.5 | 0.049 | 0.037 | 0.654 | 0.572 | 0.412 | 0.183 | 0.492 | 0.625 | 0.530 | 0.702 |
| weak | 0.5 | 0.5 | 0.032 | 0.018 | 0.380 | 0.234 | 0.339 | 0.087 | 0.472 | 0.573 | 0.547 | 0.709 |
| weak | 0.5 | 1 | 0.037 | 0.026 | 0.497 | 0.387 | 0.382 | 0.181 | 0.480 | 0.560 | 0.535 | 0.664 |
| weak | 0.5 | 1.5 | 0.052 | 0.040 | 0.718 | 0.645 | 0.414 | 0.203 | 0.508 | 0.623 | 0.540 | 0.692 |
| weak | 0.8 | 0.5 | 0.037 | 0.023 | 0.475 | 0.344 | 0.434 | 0.144 | 0.412 | 0.576 | 0.473 | 0.688 |
| weak | 0.8 | 1 | 0.044 | 0.035 | 0.590 | 0.511 | 0.437 | 0.263 | 0.444 | 0.527 | 0.492 | 0.611 |
| weak | 0.8 | 1.5 | 0.053 | 0.051 | 0.759 | 0.785 | 0.432 | 0.292 | 0.496 | 0.557 | 0.525 | 0.617 |
| strong | 0.3 | 0.5 | 0.006 | 0.003 | 0.302 | 0.163 | 0.430 | 0.035 | 0.126 | 0.299 | 0.161 | 0.352 |
| strong | 0.3 | 1 | 0.007 | 0.005 | 0.406 | 0.312 | 0.465 | 0.094 | 0.221 | 0.380 | 0.246 | 0.422 |
| strong | 0.3 | 1.5 | 0.009 | 0.007 | 0.563 | 0.530 | 0.532 | 0.209 | 0.250 | 0.401 | 0.268 | 0.432 |
| strong | 0.5 | 0.5 | 0.006 | 0.004 | 0.359 | 0.223 | 0.477 | 0.055 | 0.124 | 0.341 | 0.156 | 0.389 |
| strong | 0.5 | 1 | 0.008 | 0.005 | 0.445 | 0.371 | 0.481 | 0.141 | 0.231 | 0.379 | 0.254 | 0.417 |
| strong | 0.5 | 1.5 | 0.010 | 0.008 | 0.640 | 0.595 | 0.540 | 0.240 | 0.270 | 0.405 | 0.285 | 0.433 |
| strong | 0.8 | 0.5 | 0.007 | 0.005 | 0.395 | 0.328 | 0.514 | 0.164 | 0.155 | 0.331 | 0.181 | 0.371 |
| strong | 0.8 | 1 | 0.008 | 0.007 | 0.514 | 0.465 | 0.515 | 0.268 | 0.250 | 0.339 | 0.269 | 0.371 |
| strong | 0.8 | 1.5 | 0.011 | 0.009 | 0.747 | 0.707 | 0.548 | 0.303 | 0.298 | 0.388 | 0.310 | 0.413 |

**Table 2.**
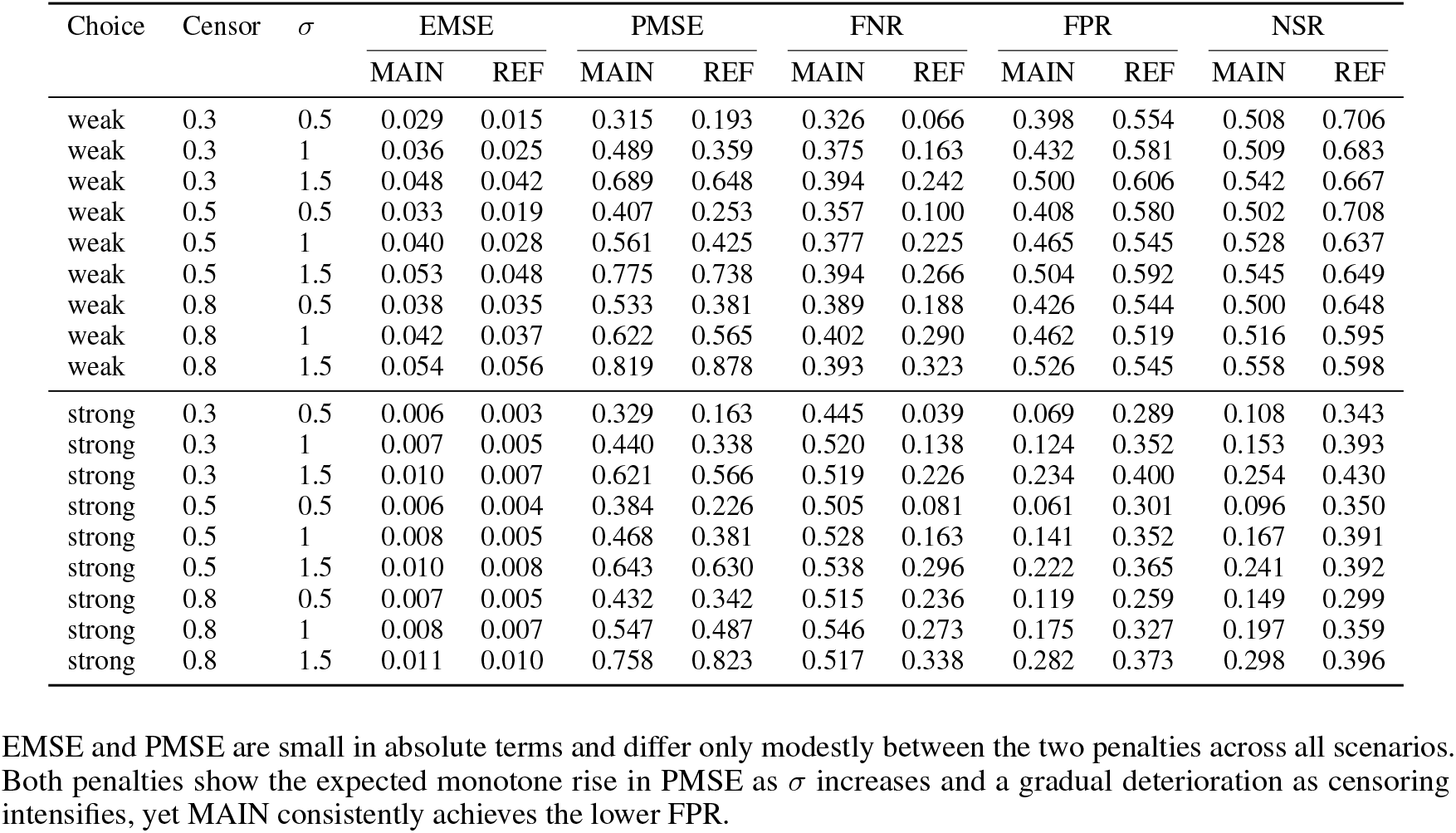
nonOverlap-simulation benchmark (100 replicates, 5-fold CV).

**Table 3.**
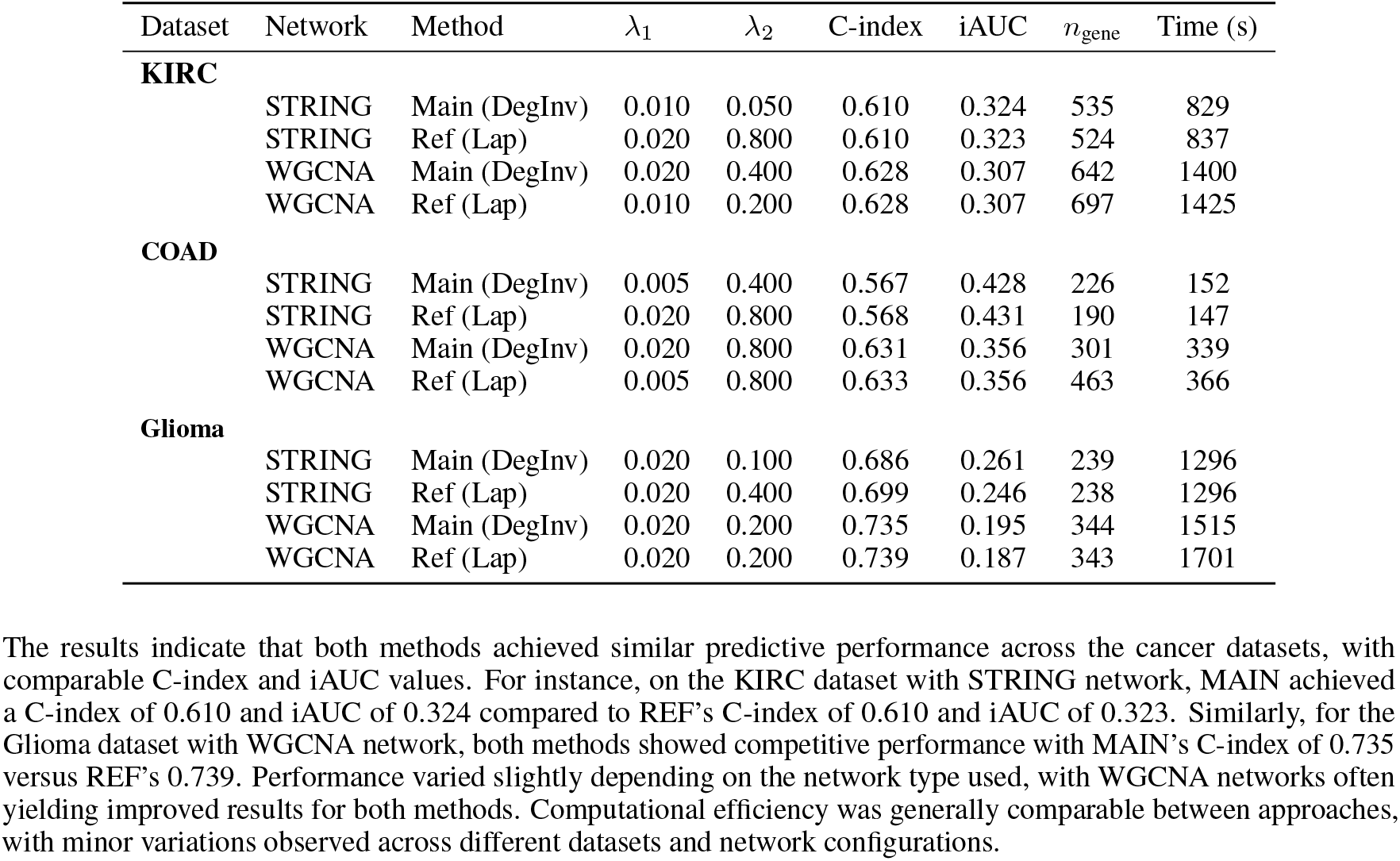
Real data application results on three cancer datasets. *λ*_1_ and *λ*_2_ are the tuning parameters; *n*_gene_ is the number of selected genes.

| Dataset | Network | Method | $\lambda_1$ | $\lambda_2$ | C-index | iAUC | $n_{\text{gene}}$ | Time (s) |
| --- | --- | --- | --- | --- | --- | --- | --- | --- |
| **KIRC** |  |  |  |  |  |  |  |  |
|  | STRING | Main (DegInv) | 0.010 | 0.050 | 0.610 | 0.324 | 535 | 829 |
|  | STRING | Ref (Lap) | 0.020 | 0.800 | 0.610 | 0.323 | 524 | 837 |
|  | WGCNA | Main (DegInv) | 0.020 | 0.400 | 0.628 | 0.307 | 642 | 1400 |
|  | WGCNA | Ref (Lap) | 0.010 | 0.200 | 0.628 | 0.307 | 697 | 1425 |
| **COAD** |  |  |  |  |  |  |  |  |
|  | STRING | Main (DegInv) | 0.005 | 0.400 | 0.567 | 0.428 | 226 | 152 |
|  | STRING | Ref (Lap) | 0.020 | 0.800 | 0.568 | 0.431 | 190 | 147 |
|  | WGCNA | Main (DegInv) | 0.020 | 0.800 | 0.631 | 0.356 | 301 | 339 |
|  | WGCNA | Ref (Lap) | 0.005 | 0.800 | 0.633 | 0.356 | 463 | 366 |
| **Glioma** |  |  |  |  |  |  |  |  |
|  | STRING | Main (DegInv) | 0.020 | 0.100 | 0.686 | 0.261 | 239 | 1296 |
|  | STRING | Ref (Lap) | 0.020 | 0.400 | 0.699 | 0.246 | 238 | 1296 |
|  | WGCNA | Main (DegInv) | 0.020 | 0.200 | 0.735 | 0.195 | 344 | 1515 |
|  | WGCNA | Ref (Lap) | 0.020 | 0.200 | 0.739 | 0.187 | 343 | 1701 |

The results indicate that both methods achieved similar predictive performance across the cancer datasets, with comparable C-index and iAUC values. For instance, on the KIRC dataset with STRING network, MAIN achieved a C-index of 0.610 and iAUC of 0.324 compared to REF’s C-index of 0.610 and iAUC of 0.323. Similarly, for the Glioma dataset with WGCNA network, both methods showed competitive performance with MAIN’s C-index of 0.735 versus REF’s 0.739. Performance varied slightly depending on the network type used, with WGCNA networks often yielding improved results for both methods. Computational efficiency was generally comparable between approaches, with minor variations observed across different datasets and network configurations.

### 3.3 Pathway Enrichment Results

The MAIN method with the STRING network and the REF method with the same network both highlight three tightly linked metabolic pathways, mTORC1 regulation by amino acids, fatty-acid *β*-oxidation and iron uptake/transport in clear-cell renal cell carcinoma, each with false-discovery rates below 0.01 (Supplementary Table 2). REF attains slightly smaller q-values yet captures nothing beyond the biology already recovered by MAIN; when the WGCNA network is used the pattern persists, with the MAIN list 3 and REF list 4 again isolating the same metabolic axis. Because disrupted lipid storage, amino-acid sensing and iron handling are core features of this tumor type (Qiu et al., 2015; Tan et al., 2022), both penalties select biologically meaningful markers, and the marginal statistical edge for REF does not translate into additional insight.

In colorectal adenocarcinoma, MAIN and REF behave almost identically on the STRING backbone, each singling out cytokine signaling, a GTSE1-centred G2/M checkpoint module and Dectin-1 signaling with comparable significance. With the WGCNA graph, however, the MAIN list 3 contains ten pathways centered on inflammation, cell-cycle regulation and proteasome function, processes firmly implicated in colorectal cancer progression (Braumüller et al., 2022; Cui et al., 2024), whereas the REF list 4 introduces several two-gene neurotransmitter-release cycles whose q-values hover just under 0.05 and lack a clear connection to colon cancer, suggesting statistical noise. Here the inverse-degree penalty provides the cleaner and more interpretable result.

For IDH-wildtype glioma, all four lists converge on an IL-17/chemokine-centered inflammatory with q-values between 2 ×10^−5^ and 5× 10^−4^, and the differences between MAIN and REF are confined to small shifts in p-value. Given the central role of IL-17-driven cytokine exchange in glioblastoma biology (Yeo et al., 2021; Wang et al., 2019), both penalties clearly retrieve relevant markers, and neither uncovers additional pathways beyond this dominant theme.

Taken together, the inverse-degree regularize reliably yields biologically coherent enrichment profiles. It matches the Laplacian alternative in clear-cell renal cell carcinoma and IDH-wildtype glioma and offers superior interpretability in colorectal adenocarcinoma when the network is derived from co-expression. Where the Laplacian shows modestly stronger raw significance, it does not deliver new biological content; conversely, when it introduces low-credibility terms, the inverse-degree method avoids them, reinforcing its value as the primary regularizer in the proposed pipeline.

### 3.4 R package

In support of these analyses, I have bundled the entire workflow into a self-contained R package, DegreeAFT, available at https://github.com/yliu38/DegreeAFT. The package automates STRING and WGCNA network construction, implements both Laplacian- and inverse-degree–penalized AFT models with built-in cross-validation, and ships with example datasets so users can reproduce every result in a few lines of code.

## 4 Discussion

### 4.1 Principal Findings and Methodological Contributions

This study presents the first systematic comparison of inverse-degree versus Laplacian penalties in the context of network-regularized AFT models for high-dimensional genomic survival analysis. Our comprehensive evaluation across multiple cancer datasets and simulation scenarios demonstrates that the inverse-degree penalty offers a valuable alternative to traditional Laplacian regularization, particularly when conservative feature selection and biological interpretability are prioritized.

The inverse-degree penalty consistently exhibited more conservative behavior in feature selection, achieving substantially lower false positive rates while maintaining competitive predictive performance. This finding aligns with previous observations by Veríssimo et al. (2016), who demonstrated that degree-based penalties can improve variable selection stability in Cox proportional hazards models. Our extension to AFT models represents a significant methodological advancement, as AFT models offer more intuitive interpretation of covariate effects on survival time scales compared to hazard-based approaches (Wei, 1992).

### 4.2 Network Structure and Biological Relevance

The superior performance of the inverse-degree penalty in pathway enrichment analysis, particularly evident in the colorectal adenocarcinoma dataset, suggests that this approach may better capture biologically meaningful gene-gene relationships. While both penalties identified relevant pathways such as mTORC1 regulation and fatty acid metabolism in renal cell carcinoma, consistent with established metabolic dysregulation in this cancer type (Qiu et al., 2015; Tan et al., 2023), the inverse-degree method demonstrated cleaner enrichment profiles with fewer potentially spurious associations.

This observation is particularly important given the growing recognition that gene regulatory networks exhibit scale-free properties with highly connected hub genes playing central roles in cellular processes (Barabási & Oltvai, 2004). The inverse-degree penalty’s approach of applying stronger shrinkage to highly connected nodes may better reflect the biological reality that hub genes are often involved in multiple pathways and may be more prone to false positive selection due to their extensive interactions (Li & Li, 2008).

### 4.3 Performance Across Cancer Types and Network Structures

Our analysis across three distinct cancer types, IDH-wildtype gliomas, KIRC, and COAD, reveals that the relative performance of network penalties can vary depending on the underlying biological context and network structure. The consistent identification of IL-17/chemokine signaling pathways in glioma analysis by both methods underscores the robustness of these approaches in capturing well-established biological mechanisms, as IL-17-mediated inflammation is a known driver of glioblastoma progression (Wang et al., 2019).

The differential performance observed between STRING protein-protein interaction networks and WGCNA co-expression networks highlights an important consideration for practitioners. Co-expression networks, derived from the data itself, may capture context-specific regulatory relationships that are not present in general protein interaction databases (Langfelder & Horvath, 2008; Zhang & Horvath, 2005). Our results suggest that the choice of network type should be informed by the specific biological context and research objectives, with co-expression networks potentially offering advantages when studying tissue-specific or disease-specific regulatory mechanisms.

### 4.4 Computational Considerations and Practical Implementation

The development of the DegreeAFT R package addresses a critical gap in available software tools for network-regularized survival analysis. While several packages exist for Cox proportional hazards models with network penalties (Simon et al., 2011), AFT models with network regularization have been less accessible to the broader research community. The computational efficiency observed in our implementation, achieved through proximal gradient optimization, makes these methods feasible for large-scale genomic studies typically encountered in cancer research.

The proximal gradient approach employed in our implementation offers several advantages over coordinate descent methods commonly used in penalized regression. The ability to handle non-smooth penalties while maintaining computational efficiency is particularly important for network-regularized models, where the penalty structure can be complex (Li & Li, 2008). Our implementation’s performance across datasets with varying dimensions (from hundreds to thousands of features) demonstrates the scalability required for modern genomic applications.

### 4.5 Limitations and Methodological Considerations

Several limitations of our study warrant discussion. First, our evaluation focused primarily on Weibull AFT models, and the generalizability to other AFT distributions (log-normal, log-logistic) remains to be established. While the Weibull distribution is commonly used in survival analysis due to its flexibility and interpretability (Klein & Moeschberger, 2003), different distributions may be more appropriate for specific cancer types or clinical contexts.

Second, our network construction relied on existing databases (STRING) and standard co-expression analysis (WGCNA), which may not capture all relevant biological relationships. Recent advances in single-cell genomics and multi-omics integration offer opportunities to construct more comprehensive and context-specific networks (Stuart & Satija, 2019). Future work should explore the integration of these emerging data types into network-regularized survival models.

The simulation study, while comprehensive, employed relatively simple network structures compared to the complexity of real biological networks. Real gene regulatory networks exhibit hierarchical organization, overlapping modules, and dynamic rewiring across conditions (Fortunato, 2010). More sophisticated simulation frameworks incorporating these features would provide additional insights into method performance under realistic conditions.

### 4.6 Future Directions

Several avenues for future research emerge from this work. The integration of multi-omics data (genomics, proteomics, metabolomics) into network-regularized survival models represents a promising direction for improving predictive accuracy and biological insight (Hasin et al., 2017). The incorporation of dynamic network information, reflecting changes in gene regulatory relationships across disease stages or treatment responses, could further enhance model performance.

The development of ensemble methods combining multiple network penalties or integrating network-regularized models with other machine learning approaches (random survival forests, neural networks) may yield improved predictive performance while maintaining interpretability (Ishwaran et al., 2008). Additionally, the extension of these methods to competing risks scenarios, where patients may experience multiple types of events, would broaden their applicability in clinical research.

The growing availability of real-world evidence from electronic health records and cancer registries provides opportunities to validate network-regularized survival models in larger, more diverse patient populations. Such validation studies are essential for translating these methodological advances into clinical practice and ensuring their robustness across different healthcare settings and patient demographics.

### 4.7 Conclusion

This study demonstrates that network-regularized AFT models with inverse-degree penalties offer a valuable tool for biomarker discovery in high-dimensional genomic survival analysis. The conservative feature selection behavior and superior biological interpretability of the inverse-degree penalty, combined with competitive predictive performance, support its adoption as a primary regularization strategy in genomic survival studies. The DegreeAFT R package provides the research community with accessible tools for implementing these methods, facilitating broader adoption and further methodological development.

The identification of prognostic gene signatures in IDH-wildtype gliomas and the demonstration of method performance across multiple cancer types underscore the translational potential of these approaches. As genomic technologies continue to advance and larger datasets become available, network-regularized survival models will likely play an increasingly important role in precision oncology and personalized treatment decision-making.

## 5.1 Weibull AFT density and survival functions

Let *T*_*i*_ be a non-negative survival time with covariate vector **x**_*i*_ ∈ ℝ^*p*^. In AFT form

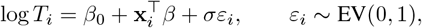

so the scale parameter is 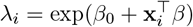 and the shape parameter is *k* = 1*/σ*.

The Weibull pdf and survival function for subject *i* are

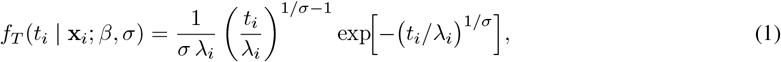

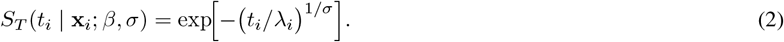

## 5.2 Penalized log-likelihood

With *δ*_*i*_ ∈ {0, 1} the event indicator, the penalized objective is

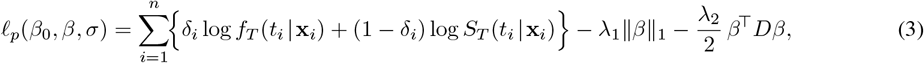

where *D* = *diag*(*B*^⊤^*B*) is the *DEGREE* matrix.

## 5.3 Log-likelihood in log-time coordinates

Define *y*_*i*_ = log *t*_*i*_ and the rescaled residual

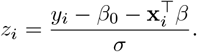

Using 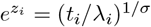, the per-observation log terms are

**Uncensored** (*δ*_*i*_ = 1):

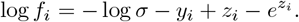

**Censored** (*δ*_*i*_ = 0):

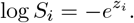

Hence

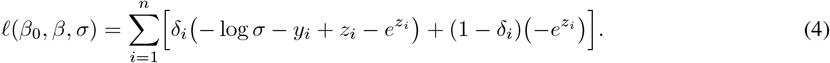

## 5.4 Gradient derivations

Write **x**_*i*_ for the *i*-th covariate vector.

**With respect to** *β*

Because *∂z*_*i*_*/∂β* = −**x**_*i*_*/σ*,

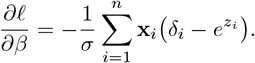

**With respect to** *β*_0_

Since *∂z*_*i*_*/∂β*_0_ = −1*/σ*,

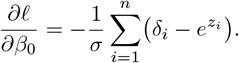

**With respect to** *σ*

Using *∂z*_*i*_*/∂σ* = −*z*_*i*_*/σ* and 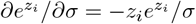,

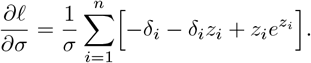

## 5.5 Optimization with Proximal Gradient Descent

Split (3) into a smooth part 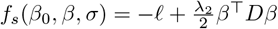 and a non-smooth part *g*(*β*) = *λ*_1_∥*β*∥_1_. For step size *η*,

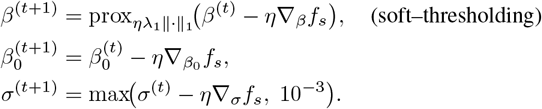

Here

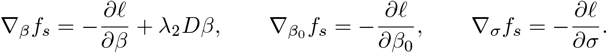

